# Biological Potencial of *Colletotrichum typhae* H.C Greene mycoherbicide for *Typha domingensis* Pers

**DOI:** 10.1101/502526

**Authors:** Cláudio Belmino Maia, Paulo Alexandre Fernandes Rodrigues de Melo, Robert Weingart Barreto, Luiz Antônio Maffia, Kedma Maria Silva Pinto, Antonia Alice Costa Rodrigues, Ilka Márcia Ribeiro de Souza Serra, Mário Luiz Ribeiro Mesquita, Janaina Marques Mondego, Aline Priscilla Gomes da Silva

**Affiliations:** Universidade Estadual do Maranhão, Centro de Ciências Agrárias, Programa de Pós-Graduação em Agroecologia, CP 65.055-098, São Luís, MA, Brazil.; Universidade Federal de Viçosa, Centro de Ciências Agrárias, Departamento de Fitopatologia.; Universidade Federal Rural de Pernambuco, Programa de Pós-Graduação em Produção Agrícola.; Universidade Estadual do Maranhão, Programa de Pós-Graduação em Agricultura e Ambiente.; Michigan State University, Department of Horticulture, East Lansing, MI 48824, United States.

**Keywords:** Biological control, Aquatic macrophytes, Leaf wetness

## Abstract

The anthropic interference in aquatic ecosystems, favors the disordered colonization of *T. domingensis*, damaging the production of hydroelectric power and river traffic. Thus, the objective of this study was to evaluate the potential of *C. typhae* as a mycoherbicide in the control of *T. domingensis*, in vitro and in greenhouse. 107 samples of symptomatic *T. domingensis* leaves were collected in flooded areas of rivers in Brazil, with identification and isolation of the collected fungal species. The concentration of inoculum was determined to evaluate the incidence and severity of the disease, the influence of temperature on mycelial growth and conidia germination, the effect of temperature and leaf wetness period on *T. domingensis* infection by *C. typhae* and the host range test. The growth of the colonies of *C. typhae* was higher at 25 to 30 ºC, there was no interference of the photoperiod on germination of the spores, but the highest percentage of germination occurred at 17.39 ºC. The influence of environmental conditions on infection of inoculated leaves of *T. dominguensis* indicated that at 15 °C and the period of leaf wetness of 48 hours promoted the highest incidence of the disease, as well as the severity for the same period of leaf wetness. The specificity test showed that *C. typhae* is specific and pathogenic to *T. domingensis*. Being this the first report of the occurrence of this pathogen in aquatic macrophytes of this species and in *T. domingensis* in Brazil.

## Introduction

*Typha domingensis* Pers. is an invasive macrophyte found in the Americas, Europe, Africa, Asia and Oceania, being considered as a native species of South America, occurring throughout Brazil [1]. It is propagated either by seeds or vegetatively, by rhizomes, with vigorous growth by the decomposition and assimilation of organic matter as a source of nutrients, reaching about seven tons of rhizomes per hectare [2]. Thus, *T. domingensis* is used as a biological filter for urban sewage, industrial effluents rich in heavy metals and erosion control in drainage channels and reservoir banks [3].

However, anthropic interference in aquatic ecosystems favors the colonization of *T. domingensis*, which may hinder the production of hydroelectric power, river traffic and agricultural irrigation [4]. In the United States, this macrophyte accounts for the degradation of almost 12.000 ha^-1^ of Florida marshes, due its aggressive growth in response to eutrophication by nitrates and phosphates from agricultural and urban waste and frequent fires [5]. In Brazil, it is estimated that the intense growth of *T. domingensis* in reservoirs of the country’s hydroelectric dams extends to about 300 ha^-1^ in the water mirror. These changes contribute to the reduction of water quality and biodiversity patterns in environments colonized by this species [6].

Up to a certain limit, the development of aquatic vegetation can be considered harmful in several ecosystems [7]. In order to reduce the environmental, social and economic impacts caused by hydrophytic plants, mechanical and biological control have been used [8,9]. The use of chemical herbicides is another option to control aquatic weed macrophytes, being allowed in countries like the United States, although its application is controversial in European countries and in Brazil [10]. This is due to the low acceptance by society, due to the excessive use of toxic agrochemicals to different plant species, besides the low number of registered products [11].

Thus, the use of weed control methods with a higher degree of specificity that reach only the target species constitutes viable alternatives [12]. Among these, the mass production of microorganisms, intended for the formulation of mycoherbicides, has been shown to be effective in weed management in several parts of the world [13]. Research results have shown the efficacy of several commercial mycoherbicides made with fungi of the genus *Colletotrichum*, with control rates reaching 90%, such as the use of *Colletotrichum truncatum* (Schwein) in *Sesbania exaltata* (Raf.) Rydb. ex AW Hill., *Colletotrichum acutatum* (Sim.) in *Hakea sericea* Schrad. and *Colletotrichum gloeosporioides* Penz. (Sacc.) in the control of *Aeschynomene virginica* L. [11].

Research on microorganisms in the prospection of mycoherbicides aims to establish optimal culture conditions for mass and durable production of the inoculum in artificial culture [14]. The pathogen should be genetically stable and specific in order to generate rapid and high disease level, with consequent death or suppression of the target plant, and should not present pathogenicity to crops of agricultural interest [15]. Thus, it is necessary to study the pathogen and its interaction with the host, as well as the conditions that predispose the plant to the pathogen, since climate variables such as temperature may influence both infection and colonization of the pathogen [16]. In addition, host specificity and preference has been a criterion used in research with *Colletotrichum* species for the biological control of plants. Some species of these pathogens are capable of infecting single hosts and, conversely, there are also *Colletotrichum* species capable of infecting multiple host species [17].

In recent years, little has been studied about these parasitic relationships between fungi and aquatic weed macrophytes, however, satisfactory results were obtained in studies with the *Eichhornia crassipes* Mart. Solms and *Cercospora rodmanni* [11]. Another important result was reported by Pitelli et al. [8], showing that fungi of the genus *Colletotrichum* have been receiving attention as potential mycoherbicides, suggesting that these pathogens have specific enzymes that promote infection and degradation of the plant cell wall. Thus, the objective of this study was to evaluate the potential of *C. typhae* as a mycoherbicide in the control of *T. domingensis*, in vitro and in greenhouse.

## Materials and Methods

### Obtainment and identification of *T. dominguensis* pathogenic isolates

A total of 107 leaf samples from symptomatic *T. domingensis* plants with necrotic spots were collected from flooded areas of the São Francisco and Doce rivers, Brazil, from municipalities of the states of Sergipe (SE), Espirito Santo (ES), Minas Gerais (MG) and Bahia (BA) (Table 1). The collected material was packed in polyethylene bags and taken to the Plant Diseases Laboratory, Department of Plant Pathology, Federal University of Viçosa, Viçosa, state of Minas Gerais, Brazil.

**Table 1.** Description of the places of origin where symptomatically T. *domingensis* leaf samples were collected, geographical coordinates, climate, temperature and rainfall (Source: Climate-data. Org, 2015). Viçosa, MG, Brazil, 2018.

| Place of origin | Latitude<br>(S) | Longitude<br>(W) | *Climate | Temperature<br>(°C) | Rainfall<br>(mm) |
| --- | --- | --- | --- | --- | --- |
| Pacatuba (SE) | 10° 27' 12" | 36° 39' 05" | As | 25.5 | 1,223 |
| Brejo Grande (SE) | 10° 25' 28" | 36° 27' 44" | As | 25.3 | 1,283 |
| Propriá (SE) | 10° 12' 40" | 36° 50' 25" | Aw | 25.9 | 821 |
| Regência (ES) | 19° 23' 27" | 40° 04' 19" | Aw | 24.3 | 1,213 |
| Vila do Riacho (ES) | 19° 49' 12" | 40° 16' 22" | Aw | 24.5 | 1,178 |
| Barra do Riacho (ES) | 19° 49' 24" | 40° 04' 20" | Aw | 24.5 | 1,162 |
| Santa Cruz do Escalvado (MG) | 20° 14' 09" | 42° 48' 50" | Aw | 22.1 | 1,146 |
| Rio Doce (MG) | 20° 14' 41" | 42° 53' 45" | Aw | 22.2 | 1,143 |
| Ipatinga (MG) | 19° 42' 23" | 42° 34' 33" | Aw | 23.8 | 1,143 |
| São Francisco (MG) | 15° 56' 55" | 45° 51' 52" | Aw | 23.6 | 1,053 |
| Juazeiro (BA) | 09° 24' 50" | 40° 30' 10" | BSh | 24.8 | 422 |
| Paulo Afonso (BA) | 09° 24' 22" | 38° 12' 53" | BSh | 25.8 | 540 |
| Xingozinho (BA) | 09° 32' 30" | 38° 01' 59" | BSh | 25.8 | 540 |

\*Classification according to [18]. http://pt.climate-data.org/country/114/

As and Aw (tropical savannah climate with dry winter season), with average temperature in any month of the year exceeding 18 °C. Winter is dry, with average rainfall less than 60 mm in at least one of the months of this season. BSh (hot semi-arid climate), with average temperature of the hottest month of the year 27.8 ° C. With a dry season during the year, with average rainfall in the driest month around 8 mm. Most of the rainfall occurs in March with an average of 83 mm.

Isolates of the collected samples were made and the fragments containing the fungal structures were grown in Petri plates containing Potato Carrot Agar (PCA) culture medium. The isolates were incubated at 25 ± 3 °C in the dark for eight days [19]. The identification of organisms at the genus level was carried out using the identification key of Barnett and Hunter [20], by observations of the structures under a stereoscopic microscope. In order to confirm the pathogenicity of the isolates, they were inoculated in *T. dominguensis* disease free seedlings produced in a greenhouse, reproducing the symptoms verified in the field. For this, fungal suspensions (2.5×10^6^ conidia / mL), plus Tween 80 (0.05%) were used and the inoculation was done by brushing. The plants were then kept in a fog chamber at 25 ºC for 48 hours. After that time, they remained in the greenhouse for the evaluation, which was carried out daily, for 30 days.

### Concentrations of *Colletotrichum* sp. to evaluate the incidence and severity of the disease

Due to the rapid sporulation in culture medium and the severity of the disease, which was assessed by fast evolution of the symptoms in the inoculated plant, only the cultures of the genus *Colletotrichum* sp. obtained in the detection test were used. To this end, the isolates were grown in potato dextrose agar (PDA) culture medium and incubated at 25 ± 1 °C in the dark for eight days. The identification of the *Colletotrichum typhae* species was carried out under a stereoscopic microscope [21], being this the first report of the occurrence of this pathogen in aquatic macrophytes of this species and in *T. domingensis* in Brazil. The sample was subcultured three times to obtain a pure culture and stored at 5 ± 1 °C after characterization of the species for further testing.

Four groups of adult and young plants of *T. domingensis*, arranged in three replicates, were used to evaluate the severity of the disease, with each replication consisting of a pot with three plants. The leaves were inoculated by brushing the conidial suspensions with 0.05% Tween 80 fixative, adjusted at four different concentrations (one for each group of plants): 2.5 x 10^4^, 2.5 x 10^5^, 2.5 x 10^6^ and 2.5 x 10^7^ conidia / mL ^-1^. The experiment was conducted in a humid chamber at 25 ± 1 °C at 90% relative humidity and transferred to a greenhouse after 48 hours. The evaluations were carried out daily, during eight days. Since there is no descriptive or diagrammatic scale of the disease, the severity was evaluated by the percentage of leaf area affected by the symptoms, which was quantified from photocopied detached symptom leaves [22] using the software Severity.exe., where the injured area was calculated, thus it was possible to correlate the symptomatology and leaf morphology with damages caused by the anthracnose disease caused by *C. typhae*.

### Influence of temperature on mycelial growth and germination of *C. typhae* conidia

For evaluation of mycelial growth, *C. typhae* mycelium discs (RWB99) from colonies with eight days of incubation in Vegetable Broth Agar (VBA) medium were placed in Petri plates, containing the same medium, one disc per plate, with five replicates with the experimental unit consisting of two Petri plates incubated at temperatures of 15, 20, 25, 30 and 35 °C in the dark. The evaluations were performed daily by measuring the diameter of the colonies in two perpendicular directions, during four days and from the averages obtained it was calculated the area below the mycelial growth curve.

To evaluate the conidia germination, a 2.5×10^6^ conidia / mL suspension of *C. typhae* was used. Four aliquots of 50μL were deposited at equidistant points in Petri plates containing agar-water medium scattered with a Drigalski loop. Five replicates were used, each containing two plates. The plates were incubated for eight days under the same conditions of the tests for evaluation of mycelial growth. After this time, aliquots of sterile distilled water were added to each plate. A fungal suspension was prepared by scraping the colonies and 100 spores of each plate were counted with the aid of a hemacytometer and an optical microscope, quantifying, in percentage, how many of these spores were germinated.

### Influence of temperature and leaf wetness period on *T. domingensis* infection by *C. typhae*

*T. domingensis* seedlings were prepared in polyethylene pots with a three liter capacity using coarse sand (3.0 μm), organic substrate and Red Latosol (1:1:1; v/v) as substrate. Four groups of fifteen plants of different ages were inoculated with a suspension of 2.5 x 10^6^ spores / ml by atomization until run-off with P-600 compressor with the “PULVERJET” P-110 pistol. After inoculation the plants were kept in a humid chamber (wire-frame vessels wrapped with internally moistened plastic) within growth chambers with the following temperatures: 15, 20, 25 and 30 ± 2 °C. Each group consisted of five subgroups of three plants, which were gradually removed from the wraps after 8, 12, 24 and 48 hours of leaf wetness. A subgroup for each treatment was placed in each growth chamber without being wrapped in plastic (corresponding to zero hour of leaf wetness). After completing 48 hours, all plants were taken to the greenhouse to evaluate the incidence and severity of the disease. Six evaluations were performed at two-day intervals, with the first evaluation on the eighth day after inoculation.

### *C. typhae* host range

In order to verify the host range of the isolate under study, inoculations were carried out on the species selected according to the phylogenetic centrifugal method [23], with modifications. Due to difficulties of obtaining other plants of the same genus and restrictions of the distribution of the Typhaceae family or even of the Typhales order, made it difficult to perform more complete analyzes for this pathogen. Thus, *C. typhae* inoculation was performed in *T. domingensis* and plants of the same order and related orders of fruit, forest and large crop species (Anacardiaceae, Apiaceae, Araceae, Arecaceae, Asteraceae, Bromeliaceae, Cannaceae, Caricaceae, Chenopodiaceae, Commelinaceae, Cyperaceae, Euphorbiaceae, Euriocaulaceae, Fabaceae, Halaceae, Laceaceae, Malaceae, Mayaceae, Maraceaeae, Musaceae, Myraceceae, Poaceae, Pontederiaceae, Rosaceae, Rubiaceae, Rutaceae, Solanaceae, Sparganiaceae, Strelitzaceae, Typhaceae, Vitaceae, Xyridaceae and Zingiberaceae). Forest species and agricultural crops were included in the evaluation, whether or not related to the target plant, and plants of species susceptible to fungi of the genus *Colletotrichum* sp., totaling 53 species. Ten leaves of each plant species were inoculated by the 2.5 x 10^6^ conidia / mL ^-1^ suspension by the brushstroke method and were kept in a humid chamber at 25 ± 1 °C with 90% relative humidity for 48 hours. After this time, they were transferred to a greenhouse and the evaluations to detect the presence or absence of symptoms caused by *C. typhae* were carried out at the 3rd, 7th and 21st days.

### Statistical analysis

The tests performed to identify the effect of temperature on mycelial growth and germination of *C. typhae* spores were conducted in a randomized complete design with five replicates, with two Petri plates each. To evaluate the influence of leaf wetness and temperature on *T. dominguensis* infection by *C. typhae*, the trials were conducted in a randomized complete design with 20 treatments and three replicates, each replicate consisting of a pot with three plants. The statistical editor used for the analyzes was SAEG.

The effect of the temperatures was evaluated by the analysis of variance and linear regression analysis. The equations were chosen based on the significance of the regression coefficients, adopting the level of 1% of probability. In the photoperiod effect test, the treatment means were compared by the Tukey test at a significance level of 5%.

In order to evaluate the effect of temperature and leaf wetness on the incidence and severity of the disease, a experiment was conducted in a complete randomized design, with four temperatures and five leaf wetness periods, treated in a factorial scheme with two factors (temperature and leaf wetness periods) and the data submitted to a linear regression analysis.

## Results and Discussion

### Effect of different temperatures on *C. typhae* conidia germination

The microflora associated with *T. dominguensis* plants was identified as *Colletotrichum typhae* (RWB-99, CBM-24, 26, 35, 94 and 97), *Cercospora* sp. (CBM-32, 97), *P. dichota* (CBM-14, 16, 24, 26, 94, 97), *Phoma* sp. (CBM-05, 97), *C. typharum* (CBM-5, 13, 14, 16, 23, 32, 97, 24) and *Stenella* sp. (CBM-13), but only the *C. typhae* isolates were pathogenic to *T. domingensis* at the concentration used in the pathogenicity test. For this reason, the other tests were carried out only with *C. typhae*, to verify their bioherbicidal capacity on *T. dominguensis*.

After analyzing the pathogenicity of the isolate at different inoculum concentrations, a development pattern of the anthracnose symptoms was observed at the concentration of 2.5 x 10^6^ spores / ml on the eighth day after inoculation, thus, the infection started with lesions slightly visible although homogeneously distributed in the lap of the plants reaching a leaf area of 19%. In this study, levels from 0 to 100% of injured leaf area were identified, and the images obtained aided in the evaluation of the other trials.

The conidia produced by *Colletotrichum* sp. are survival structures. They are important in infection of the host and in the propagation of these pathogens. Despite its relevance to the fungal life cycle, the conidia biology has not been extensively investigated. In this study, the first description of the *C. typhae* conidia germination, whose ideal thermal condition was verified at a temperature of 20 °C, was observed, whereas the lowest spore production index was observed at the temperature of 35 °C (Fig. 1A).

**Figure 1.**
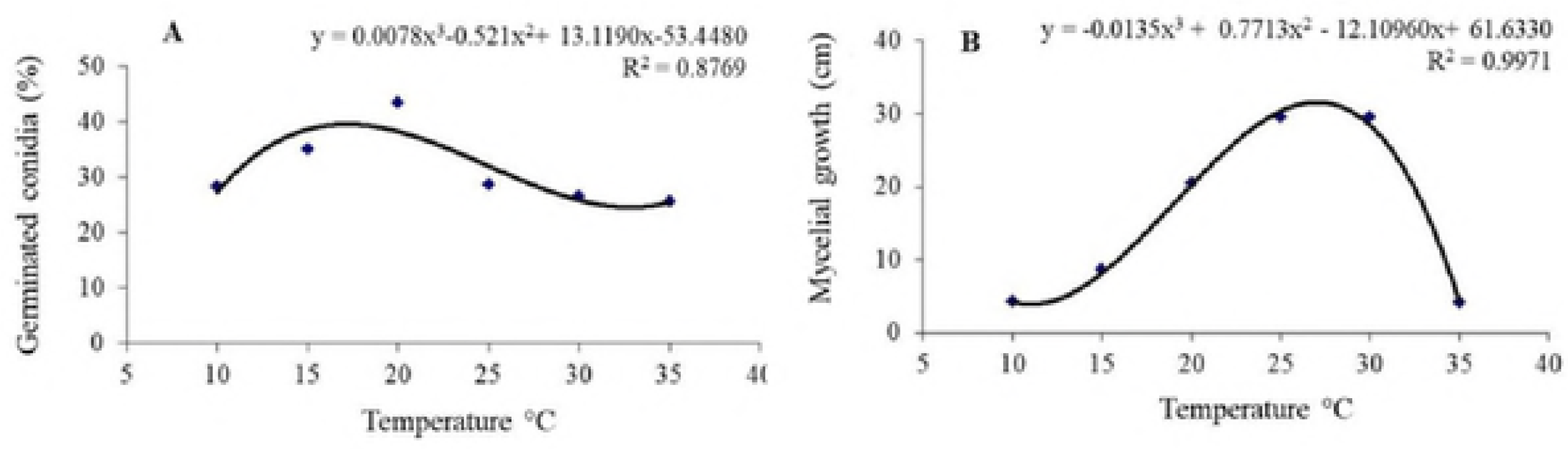
Effect of different temperatures, on *C. typhae* conidia germination in agar-water (A) and micelial growth in VBA (B).

Similar results were obtained by Estrada et al. [24] who found that the optimal temperature for sporulation of *C. gloeosporioides* was between 20 and 25 °C. However, Poltronieri et al. [25] reported that the optimal temperature for conidia germination may vary for different species of the genus *Colletotrichum* sp., Harding and Raizada [11] observed that the germination of *Colletotrichum musae* B & MAC conidia is stimulated between 27 and 30 °C while Couto and Menezes [26], demonstrated that the *Colletotrichum coccodes* Wallr. sporulation occurs between 20 and 30 °C and for *Colletotrichum lagenarium* (Pass.) Ellis & Halst. at 16 °C.

The temperature data on the disease incidence were adjusted to the cubic regression model for mycelial growth and conidial germination variables. The conidia germination and the temperature range between 25 and 30 ºC provided the highest *C. typhae* mycelial growth (Fig. 1A and 1B).

The inhibitory effect of temperature on fungi growth is variable, but most pathogens show better development at 20 to 25 °C. For *C. acutatum*, McKay et al. [27] reported that the optimum temperature for mycelial growth was 25 °C, and Harding and Raizada [11] stated that elevated temperatures around 35.5 °C paralyze *C. gloeosporioides* mycelial growth, corroborating with the results obtained in this study.

The relationship of a pathogen to the host may vary from one host to another and in this case, the interactions between this pathogenetic system are still not well understood [17]. Due to this interaction, studies on microorganisms for biological control of plants, aim to establish ideal culture conditions for mass and durable production of the inoculum in artificial culture [14]. Climate variables such as temperature, for example, influences the rate of fungal reproduction, physiological conditions of the host, growth and aggressiveness of pathogens. Thus, the knowledge of the interaction of the pathogen with environmental factors has a practical meaning, since the environment can alter its pathogenicity [28].

### Effect of leaf wetness on the incidence of the desease and severity caused by *C. typhae*

The disease incidence data, when submitted to different temperatures and leaf wetness periods, were adjusted to the linear regression model, being a decreasing incidence in relation to the increase in temperature, and increasing in relation to the leaf wetness period (Fig 2 A and B). The temperature influenced the formation of chlorotic lesions on the leaf blade surface of *T. domingensis* plants, with the exception of 20% of the plants kept at temperatures of 15, 20 and 30 °C, which had no symptoms of anthracnose in the first evaluation that occurred eight days after inoculation. It was also found that at temperatures of 15 and 20 °C, the maximum incidence of the disease was 95 and 90%, respectively (Fig 2A). On the other hand, during the evaluations of leaf wetness periods, it was observed that after the period of 48 hours, in addition to the influence of the temperature, the duration of the leaf wetness period favored the incidence of *C. typhae* with maximum percentage rate of 95% of symptomatic plants (Fig 2B).

**Figure 2.**
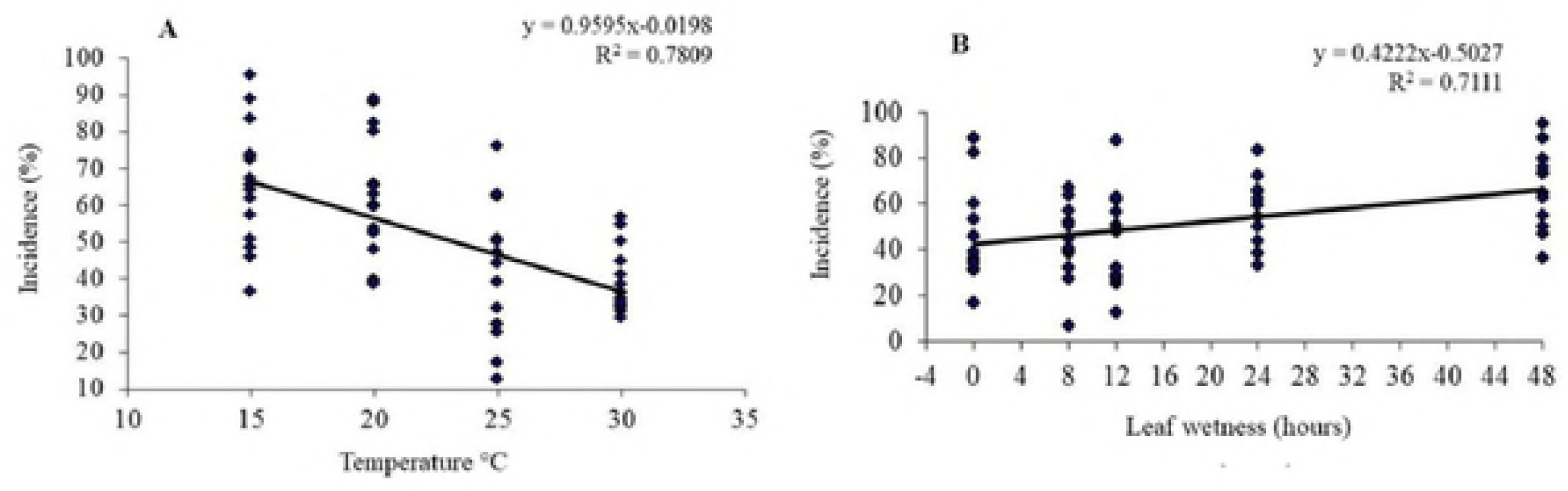
Effect of leaf wetness on the incidence of the desease caused by *C. typhae* em plants of *T. domingensis*, in the greenhouse.

With regards to the severity of the disease, it was observed that at zero hour of leaf wetness, small lesions near the lap of the plants were contacted, whereas in the higher parts of the plants these lesions were very scarce (Fig 3). Lima et al. [29] evaluating the development of phytopathogens, reported the absence of disease in zero wetness, indicating that the incidence of the fungus *Puccinia kuehnii* may be related to the need of free water on the leaves of the inoculated plants and a wetness period for at least 12 hours for fungal infection to occur. However, by the results obtained in this study, probably the presence of anthracnose symptoms in the lap of *T. domingensis* plants at zero wetness period with a rate of 85% is due to the proximity of these areas to the water in the pot, where it was possible to create a microclimate that has favored conidial germination, infection and lesion development.

**Figure 3.**
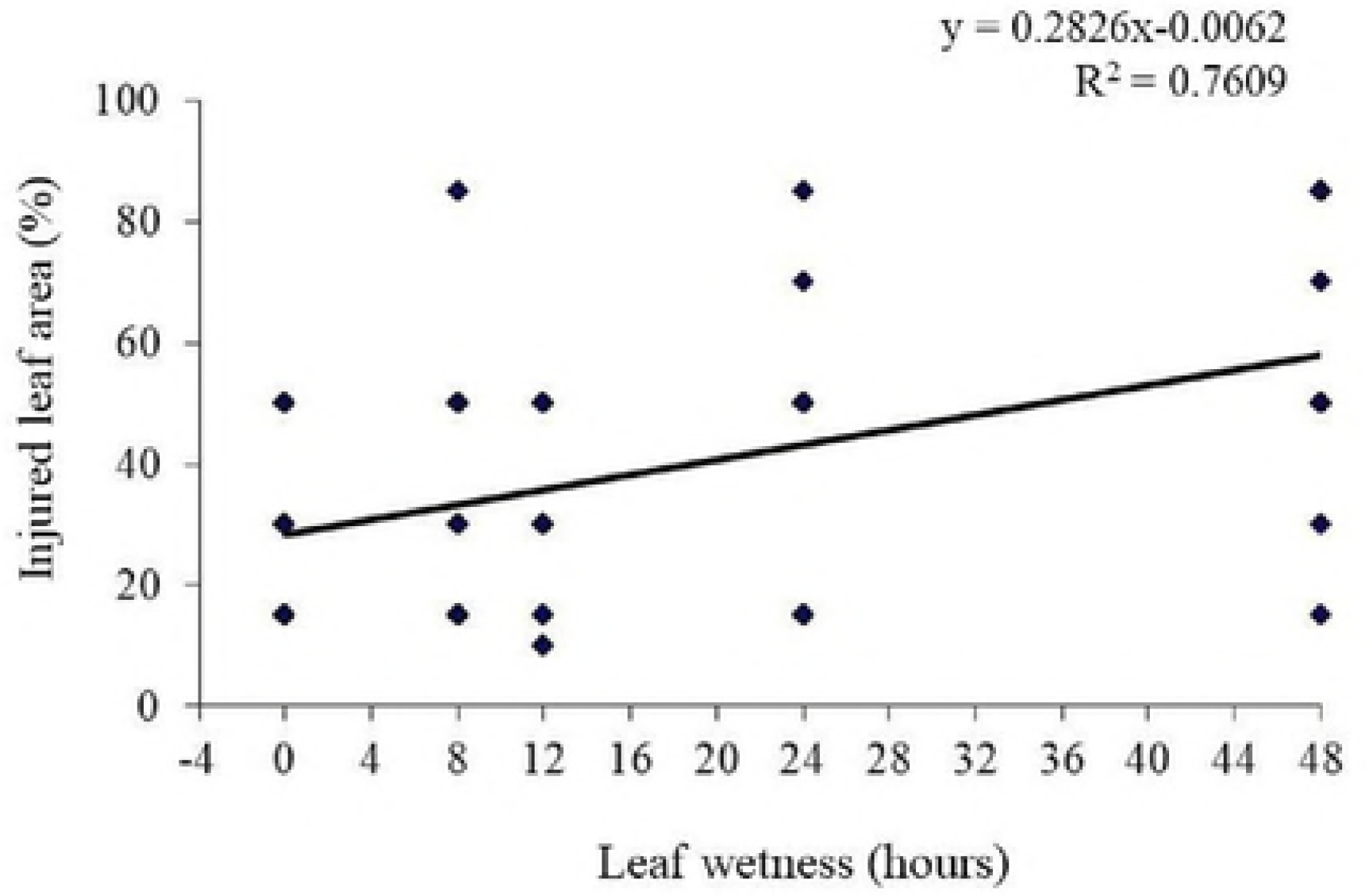
Effect of leaf wetness on the severity of the desease caused by *C. typhae* in plants of *T. domingensis*, in the greenhouse.

Spores of most pathogenic fungi require free water for hours at the host surface for germination and consequent penetration. The increase in disease intensity accompanies the wetness period up to a certain threshold, from which the maximum amount of disease is reached and it is no longer influenced by additional wetness periods [30]. According to Lima et al. [29], from a certain concentration of infection points, lesions that form first, may hinder the development of others that result from subsequent infections. However, in the present study the highest progression of both disease incidence and severity was detected with increasing leaf wetness time duration (Figs 2B and 3).

The highest severity of symptoms detected was observed in 8, 24 and 48 hours leaf wetness periods, with rates of 82, 85 and 88%, respectively. There is little research on the epidemiology of anthracnose caused by *C. typhae*; in addition, the results obtained in this study are the first report of this disease in *T. domingensis* plants, so further studies are probably needed to confirm these results (Fig 3).

Climate variables such as temperature and precipitation influence the development of anthracnose symptoms. Under high temperature conditions, pathogens can grow rapidly and severe outbreaks of disease are expected in environments with high relative humidity [31]. According to Soares-Colletti and Lourenço [32], the infection rate is high in the range of 15 to 30 ºC for *C. gloeosporioides*, with the ideal temperature around 25 ºC, however, for *C. acutatum* as for *C. typhae* a maximum incidence of this pathogen occurred at 20 °C. Regarding the wetness period, the results obtained in this study with *C. typhae* coincide with those found by Soares, Lourenço and Amorim [33], who reported that the greater severity of anthracnose caused by *C. acutatum* and *C. gloeosporioides* was high after 48 hours.

From the results obtained in this study during the evaluation of the incidence and severity of anthracnose in plants of *T. domingensis*, it was verified that the amount of hours of the wetness period needs to be continuous reaching its peak after 48 hours. In order to cause infection in plants, *C. typhae* probably requires several short wetness rather than a single long period. Among the factors that have most restricted the good performance of potential mycoherbicides is the need for a long duration of the leaf wetness period for the infection to occur [34]. However, these restrictions can be overcome by methods of artificial inoculation, since the moisture level and inoculation during the leaf wetness period can be increased by the more efficient use of wetness agents or emulsions, or by applying the fungus to gel or by using granular preparations [35,36].

Most of the studied potential plant mycoherbicides have been based on formulations of fungi species. The biological control that is involved in the application of fungal spores propagation in concentrations that do not occur in nature, is the most used strategy in the use of mycoherbicides based on these pathogens [37]. Several studies indicate that fungi of the genus *Colletotrichum* receive attention as potential mycoherbicides, since they have enzymes that degrade the plant cell walls, suggesting that some of these proteins may have specific roles in plant infection. There is also evidence that both species of *Colletotrichum* have the ability to produce indole-acetic acid, whose derivatives are well-established models of herbicides [38].

However, anthracnose caused by species of the genus *Colletotrichum* is a common and destructive disease in diverse agricultural crops and forest species. Its global occurrence causes significant losses in tropical and subtropical regions [39]. Although the genus contains species with different lifestyles, plants can be infected during any stage of development and symptoms appear after colonization of fungi, which is characterized by necrotic lesions in leaves [40]. Therefore, due to the lack of information about its occurrence and aggressiveness, there is a need to determine the susceptibility of plant species to validate the prospection of mycoherbicides, adopting the pathogenicity test proposed by Wapshere [23], widely used in biological control programs.

### Pathogenicity of the Colletotrichum hyphae isolate inoculated at room temperature

In this study, the host range was evaluated using 53 plants, among species and / or varieties included in the test, and only *T. domingensis* was susceptible to *C. typhae*, suggesting high specificity for the analyzed isolate (Table 2). The *C. typhae* isolate obtained in the survey produced symptoms similar to those obtained in the collection areas, with lesions on the leaves, both in the adaxial and abaxial parts, punctiform, yellow with chlorotic halo, becoming necrotic to the center with a light brown color. Based on these results, it can be inferred that this pathogen has effects mainly on the aerial part of *T. domingensis*, therefore, it is necessary to carry out more studies mainly at the molecular level, in order to confirm its mycoherbicide potential. It should be considered that a mycoherbicide may not necessarily have the same effect on plants as a chemical herbicide. However, these compounds have the potential to provide a competitive advantage for seedling growth through infection and growth retardation of weed seedlings. Therefore, isolation and structural characterization and mode of action of phytotoxins produced by pathogenic fungi for weeds, including aquatic invasive plants, should be investigated [41].

**Table 2.** Pathogenicity of the ecogenicidade Colletotrichum hyphae isolate inoculated at room temperature in humid chamber.Viçosa, MG, Brazil, 2018.

| Family | Species/Variety | Result* |
| --- | --- | --- |
| Anacardiaceae | *Mangifera indica* L./ Ubá | - |
| Apiaceae | *Daucus carota* L./Brasília | - |
| Araceae | *Pistia stratiotes* L. | - |
| Arecaceae | *Cocus nucifera* L./ Anã | - |
| Asteraceae | *Helianthus annuus* L. /UFV-10 | - |
| Bromeliaceae | *Ananas comosus* (L.) Merr./ Pérola | - |
| Cannaceae | *Cana denudata* Rosc. | - |
| Caricaceae | *Carica papaya* L. /Formosa | - |
| Chenopodiaceae | *Beta vulgaris* L./Wonder | - |
| Commelinaceae | *Commelina bengalensis* L. | - |
|  | *Tradescantia zebrina* Hort. ex Bosse. | - |
|  | *Tradescantia pallida* (Rose) Hunt. | - |
|  | *Callisia repens* (Jacq) L. | - |
|  | *Commelina erecta* L. | - |
| Cyperaceae | *Cyperus esculentus* L. | - |
|  | *Cyperus rotundus* L. | - |
| Euphorbiaceae | *Manihot esculenta* Crantz. | - |
|  | *Hevea brasiliensis* Willd. | - |
| Euriocaulaceae | *Paepalanthus* sp. | - |
| Haloragaceae | *Myriophyllum aquaticum* (Vell.) Verdc | - |
| Fabaceae | *Glycine max* L. / UFV-16 | - |
|  | *Phaseolus vulgares* L. | - |
|  | *Rachis hypogaea* L. / IAC-Jimbo | - |
| Juncaceae | *Eleocharis filiculmis* Kunth. | - |
| Lauraceae | *Persea americana* Mill. / Wagner | - |
| Liliaceae | *Allium cepa* L. / Monte Alegre | - |
| Malvaceae | *Gossypium hirsutum* L. / IAPAR 4PR.1 | - |
|  | *Gossypium hirsutum* L. / IAC - 22 | - |
| Mayacaceae | *Mayaca* sp. | - |
| Marantaceae | *Marantha* sp. | - |
| Moraceae | *Ficus carica* L. / Roxo de Valinhos | - |
| Musaceae | *Musa* sp./Prata | - |
| Myrtaceae | *Psidium guajava* L. / Campos | - |
| Poaceae | *Oryza sativa* L. / BR IRGA 409 | - |
|  | *Oryza sativa* L. / CICA-8 | - |
|  | *Echinochloa polystachya* (H.B.K.) Hitch | - |
|  | *Saccharum* sp. / Cana Caiana | - |
|  | *Zea mays* L. | - |
|  | *Sorghum bicolor* (L.) Moench/ BR - 700 | - |
| Pontederiaceae | *Eichhornia azurea* (Swartz) Kunth | - |
|  | *Eichhornia crassipes* (Mart.) Solms | - |
| Rosaceae | *Prunus* sp / REDHAVEN | - |
| Rubiaceae | *Coffea arabica* L. / CATUAI - 2144 | - |
| Rutaceae | *Citrus* sp. / Taiti | - |
| Solanaceae | *Nicotiana tabacum* L. / XALTHI | - |
|  | *Lycopersicon sculentum* Mill. /Santa Clara | - |
|  | *Capsicum annuum* L. / Casca Dura | - |
| Sparganiaceae | *Sparganicum* sp. | - |
| Strelitzaceae | *Strelitzia reginae* Banks | - |
| Typhaceae | *Typha domingensis* Pers. | + |
| Vitaceae | *Vitis* sp. /Niagara Rosada | - |
| Xyridaceae | *Xyris* sp | - |
| Zingiberaceae | *Zingiber officinale* Rosc. | - |

\* (+) presence of symptoms and signs of the pathogen, (-) absence of symptomatology.

Rangel-Peraza et al. [2] considered submerged aquatic plants as the most problematic, since they drastically reduce water flow, are rapid in invasion of new areas and difficult to manage or control. The perennial growth habit and the formation of monophytic colonies seem to make them the ideal target of chemical control. However, the use of chemical herbicides is often compromised by problems related to the aquatic environment, including dilution and contact time in tap water.

### Conclusions

The growth of the *C. typhae* colonies in artificial medium is higher when they are submitted to temperatures of 25 to 30 ºC, whereas, for spore germination, there was no photoperiod interference, but the highest percentage of germination occurred at 17.39 ºC. The influence of environmental conditions on the infection in artificially inoculated leaves of *T. domingensis* indicated that the temperature of 15 ºC and 48 hours of leaf wetness promoted the highest disease incidence, as well as the severity for the same period of leaf wetness. The specificity test showed that *C. typhae* is very specific since it was pathogenic only to *T. domingensis*. Being this the first report of the occurrence of this pathogen in aquatic macrophytes of this species and in *T. domingensis* in Brazil.

## Acknowledgements

The authors acknowledge the Fundação de Amparo à Pesquisa e ao Desenvolvimento Científico e Tecnológico do Maranhão-FAPEMA.

## Author Contributions

### Conceptualization

Cláudio Belmino Maia, Robert Weingart Barreto, Luiz Antônio Maffia

### Data curation

Cláudio Belmino Maia, Paulo Alexandre Fernandes Rodrigues de Melo, Kedma Maria Silva Pinto, Ilka Márcia Ribeiro de Souza Serra, Mário Luiz Ribeiro Mesquita, Aline Priscilla Gomes da Silva.

### Funding acquisition

Luiz Antônio Maffia

### Investigation

Cláudio Belmino Maia, Paulo Alexandre Fernandes Rodrigues de Melo, Robert Weingart Barreto

### Methodology

Cláudio Belmino Maia

### Supervision

Robert Weingart Barreto, Luiz Antônio Maffia

### Validation

Cláudio Belmino Maia

### Visualization

Cláudio Belmino Maia, Paulo Alexandre Fernandes Rodrigues de Melo, Ilka Márcia Ribeiro de Souza Serra, Janaina Marques Mondego.

### Writing – original draft

Cláudio Belmino Maia, Paulo Alexandre Fernandes Rodrigues de Melo, Kedma Maria Silva Pinto

### Writing – review & editing

Cláudio Belmino Maia, Paulo Alexandre Fernandes Rodrigues de Melo, Robert Weingart Barreto, Luiz Antônio Maffia, Kedma Maria Silva, Antonia Alice Costa Rodrigues, Ilka Márcia Ribeiro de Souza Serra, Mário Luiz Ribeiro Mesquita, Janaina Marques Mondego, Aline Priscilla Gomes da Silva

